# A neuronal MAP kinase constrains growth of a *C. elegans* sensory dendrite throughout the life of the organism

**DOI:** 10.1101/316315

**Authors:** Ian G. McLachlan, Isabel Beets, Mario de Bono, Maxwell G. Heiman

## Abstract

Neurons develop elaborate morphologies that provide a model for understanding cellular architecture. By studying *C. elegans* sensory dendrites, we previously identified genes that act to promote the extension of ciliated sensory dendrites during embryogenesis. Interestingly, the nonciliated dendrite of the oxygen-sensing neuron URX is not affected by these genes, suggesting it develops through a distinct mechanism. Here, we use a visual forward genetic screen to identify mutants that affect URX dendrite morphogenesis. We find that disruption of the MAP kinase MAPK-15 or the β_H_-spectrin SMA-1 causes a phenotype opposite to what we had seen before: dendrites extend normally during embryogenesis but begin to overgrow as the animals reach adulthood, ultimately extending up to 150% of their normal length. SMA-1 is broadly expressed and acts non-cell-autonomously, while MAPK-15 is expressed in many sensory neurons including URX and acts cell-autonomously. MAPK-15 acts at the time of overgrowth, localizes at the dendrite ending, and requires its kinase activity, suggesting it acts locally in time and space to constrain dendrite growth. Finally, we find that the oxygen-sensing guanylate cyclase GCY-35, which normally localizes at the dendrite ending, is localized throughout the overgrown region, and that overgrowth can be suppressed by overexpressing GCY-35 or by genetically mimicking elevated cGMP signaling. These results suggest that overgrowth may correspond to expansion of a sensory compartment at the dendrite ending, reminiscent of the remodeling of sensory cilia or dendritic spines. Thus, in contrast to established pathways that promote dendrite growth during early development, our results reveal a distinct mechanism that constrains dendrite growth throughout the life of the animal, possibly by controlling the size of a sensory compartment at the dendrite ending.

**AUTHOR SUMMARY:** Lewis Carroll’s Alice told the Caterpillar, “Being so many different sizes in a day is very confusing.” Like Alice, the cells of our bodies face a problem in size control – they must become the right size and remain that way throughout the life of the organism. This problem is especially relevant for nerve cells (neurons), as the lengths of their elaborate dendrites determine the connections they can make. To learn how neurons control their size, we turned not to a Caterpillar but to a worm: the microscopic nematode *C. elegans*, in which single neurons can be easily visualized and the length of each dendrite is highly predictable across individuals. We focused on a single dendrite, that of the oxygen-sensing neuron URX, and we identified two genes that control its length. When these genes are disrupted, the dendrite develops correctly in embryos but then, like Alice, grows too much, eventually extending up to 1.5x its normal length. Thus, in contrast to known pathways that promote the initial growth of a dendrite early in development, our results help to explain how a neuron maintains its dendrite at a consistent length throughout the animal’s life.

## INTRODUCTION

Neurons embody the adage that structure determines function in biology. The geometries and lengths of axons and dendrites determine the path and timing of information flow. Dendrites, in particular, are sculpted in ways that control how information is integrated and computed. One well-studied aspect of dendrite morphogenesis is the shaping of subcellular compartments called dendritic spines, which are micron-scale protrusions that serve as sites of information transfer via synapses. Work from many groups has elucidated how the lengths and shapes of dendritic spines correlate with synaptic activity [1,2]. An analogous subcellular compartment, called a sensory cilium, serves as the site of information transfer on sensory receptor neurons (including photoreceptor rods and cones, olfactory neurons, and auditory hair cells) [3]. Similar to dendritic spines, sensory cilia exhibit highly regulated morphologies that can be altered by sensory input [4,5]. In addition to these very conspicuous structures, dendrites may also contain less obvious subcellular compartments that aid in signal processing; for example, excitatory and inhibitory synapses are often distributed non-uniformly along dendrites, suggesting *de facto* compartmentalization [6,7]. Further, studies of signal propagation within dendrites have suggested that dendritic branches may serve as discrete computational units [8]. Yet, we know relatively little about how subcellular regions of dendrites, other than spines and cilia, are shaped.

*C. elegans* provides a powerful model for dissecting the control of dendrite morphogenesis. It exhibits highly stereotyped neuronal anatomy, such that the position of each neuron and its pattern of neurite outgrowth, branching, and connectivity have been extensively catalogued and are highly reproducible across individuals [9]. Most *C. elegans* neurons extend one or two simple, unbranched processes that often have mixed pre- and post-synaptic identity and thus cannot be classified as strictly dendritic or axonal in character [9]. In contrast, most of its sensory neurons have a bipolar morphology with one neurite that solely receives information from the environment and is therefore a dendrite [10]. While *C. elegans* has been used extensively as a model for axon development [11], considerably less attention has been paid to its dendrites. Most of the work has focused on interesting sensory neurons with complex branched dendrites, and has revealed mechanisms of dendritic branch formation and spacing [12–23], dendritic self-avoidance [24], and even dendrite tiling [25]. By comparison, dendrites with simple unbranched morphologies have been used as a model for understanding axon-dendrite polarity [26,27]. They also offer a powerful reductionist system for identifying mechanisms that control dendrite length. Most sensory neurons in the head extend a simple unbranched dendrite to the nose and terminate in a sensory cilium that receives information from the environment [10,28,29]. Through studying one such class of neurons, the amphids, we showed that dendrite length is controlled by a process called retrograde extension, in which the neuron is born at the embryonic nose and attaches a short dendrite there, after which the cell body migrates away, stretching out its dendrite behind it [30]. The extracellular matrix protein DYF-7 is required to anchor the dendrite ending at the nose, and *dyf-7* mutants exhibit severely shortened dendrites [30].

Another sensory neuron in the head, URX, offers an interesting contrast to the amphid. URX also extends a simple unbranched dendrite to the nose but it does not terminate in a sensory cilium and its dendrite length is unaffected in a *dyf-7* mutant [10,28]. URX is an oxygen-sensing neuron that detects oxygen partly through guanylate cyclase proteins that localize to a subcellular region at the dendrite ending that is neither a sensory cilium nor a dendritic spine [31–35]. We reasoned that study of URX morphogenesis would reveal distinct mechanisms for the control of dendrite length, possibly including the organization of subcellular regions within dendrites. Therefore, we undertook a forward genetic screen to identify mutants that affect URX dendrite morphology. We identified a class of mutants that affect dendrite length in the direction opposite to what we had seen before: in these mutants, the URX dendrite dramatically overgrows up to 150% of its normal length. Analysis of one of these overgrowth mutants identified a novel cell-autonomous mechanism that constrains sensory dendrite growth in an age- and activity-dependent manner.

## RESULTS

### Isolation of URX dendrite morphogenesis mutants

The two bilaterally symmetric URX neurons are bipolar, with cell bodies situated posterior to the nerve ring, and a simple unbranched dendrite projecting to the tip of the nose (Fig. 1A) [9,10,28]. To identify genes involved in URX morphogenesis, we chemically mutagenized hermaphrodites bearing an integrated GFP reporter for URX (*flp-8*pro:GFP, expressed strongly in URX and more dimly in the AUA neuron; [36]). We visually screened non-clonal F2 progeny for individuals with defective URX dendrites. From this screen, we isolated 17 recessive mutants that affect URX morphogenesis. Eight mutants exhibit shortened dendrites with few other morphological defects and will be reported elsewhere. The remaining nine mutants were grouped into four phenotypic classes: (I) dendrite overgrowth (Fig. 1B; *hmn5*, *hmn6*, *hmn17*); (II) branching or swelling near the dendrite ending (Fig. 1C; *hmn2*); (III) pleiotropic morphological defects including misplaced cell bodies with shortened dendrites (Fig. 1D; *hmn16*) or grossly disorganized or missing dendrites (Fig. 1E; *hmn11*, *hmn14*, *hmn15*); and (IV) loss of *flp-8*pro:GFP expression in URX (Fig. 1F; *hmn13*) suggesting a defect in specifying URX cell fate. The penetrance of these phenotypes is shown in Table I.

**Table 1.**
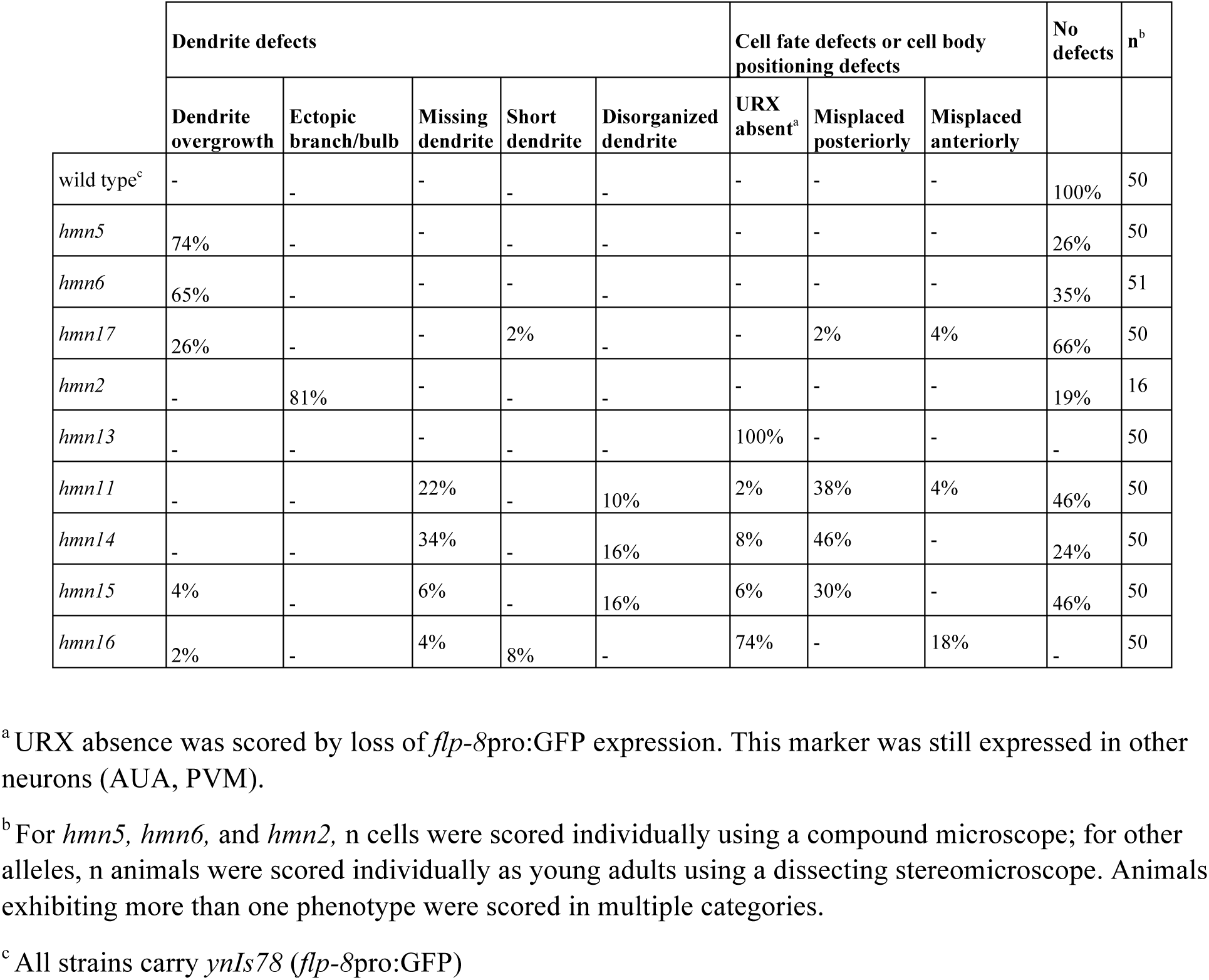
Phenotypes of URX development mutants.

|  | Dendrite defects |  |  |  |  | Cell fate defects or cell body positioning defects |  |  | No defects | n<sup>b</sup> |
| --- | --- | --- | --- | --- | --- | --- | --- | --- | --- | --- |
|  | Dendrite overgrowth | Ectopic branch/bulb | Missing dendrite | Short dendrite | Disorganized dendrite | URX absent<sup>a</sup> | Misplaced posteriorly | Misplaced anteriorly |  |  |
| wild type<sup>c</sup> | - | - | - | - | - | - | - | - | 100% | 50 |
| *hmn5* | 74% | - | - | - | - | - | - | - | 26% | 50 |
| *hmn6* | 65% | - | - | - | - | - | - | - | 35% | 51 |
| *hmn17* | 26% | - | - | 2% | - | - | 2% | 4% | 66% | 50 |
| *hmn2* | - | 81% | - | - | - | - | - | - | 19% | 16 |
| *hmn13* | - | - | - | - | - | 100% | - | - | - | 50 |
| *hmn11* | - | - | 22% | - | 10% | 2% | 38% | 4% | 46% | 50 |
| *hmn14* | - | - | 34% | - | 16% | 8% | 46% | - | 24% | 50 |
| *hmn15* | 4% | - | 6% | - | 16% | 6% | 30% | - | 46% | 50 |
| *hmn16* | 2% | - | 4% | 8% | - | 74% | - | 18% | - | 50 |

<sup>a</sup> URX absence was scored by loss of *flp-8pro:GFP* expression. This marker was still expressed in other neurons (AUA, PVM).
<sup>b</sup> For *hmn5*, *hmn6*, and *hmn2*, n cells were scored individually using a compound microscope; for other alleles, n animals were scored individually as young adults using a dissecting stereomicroscope. Animals exhibiting more than one phenotype were scored in multiple categories.
<sup>c</sup> All strains carry *ynIs78 (flp-8pro:GFP)*

**Figure 1.**
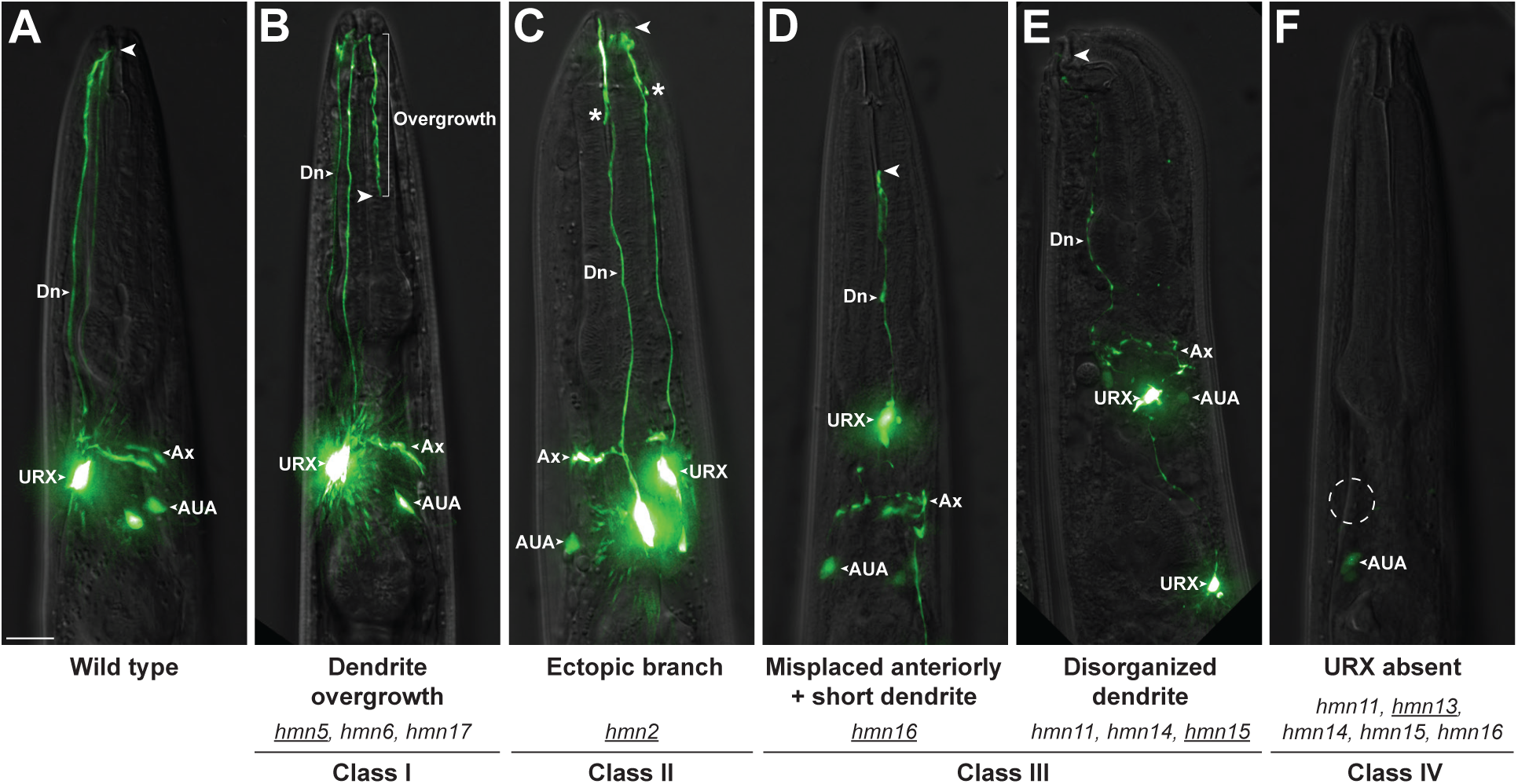
Phenotypic classes of URX dendrite morphogenesis mutants. (A) Wild-type and (B-F) mutant animals showing representative classes of phenotypes isolated in a visual forward genetic screen for URX dendrite defects. (B) Dendrite overgrowth. (C) Ectopic branch, asterisk. (D, E) Pleiotropic defects including misplaced cell bodies, shortened dendrites, and disorganized processes. (F) Loss of marker expression; dotted circle is expected position of URX. All animals express *flp-8*pro:GFP to mark URX (and, more dimly, AUA). Dn, dendrite. Ax, axon. Arrowhead, dendrite ending. Scale bar, 10 μm. Mutants exhibiting each phenotype are indicated. Mutant shown in the image is underlined. Prevalence of each phenotype is shown in Table I.

The overgrowth (Class I) mutants will be described in detail below. The branching (Class II) mutant exhibited a bulge in most URX neurons (n=13/16) at a stereotyped position along the length of the dendrite (80.4 ± 3.4% of the distance from the cell body to the dendrite ending). In 7/13 cases, this took the form of a short branch. We performed whole-genome sequencing of pooled recombinants and identified a premature termination codon in *tni-3* (Supp. Fig. 1A). A fosmid containing wild-type *tni-3* completely rescued the mutant phenotype, suggesting this is the causative mutation (Supp. Fig 1B). *tni-3* encodes one of four *C. elegans* homologs of troponin I, an inhibitory component of the troponin complex which regulates muscle contraction [37]. *tni-3* is expressed in head muscle cells [38] and the location of the ectopic dendrite branch is consistent with the URX dendrite invading into the space between these cells, suggesting that the branching phenotype may be a secondary consequence of defects in head muscle. The Class III and IV mutants were not characterized further.

### URX dendrite overgrowth mutants are alleles of *mapk-15* and *sma-1*

We chose to focus on the overgrowth mutants because the phenotype is visually striking and highly penetrant. Rather than ceasing growth at the nose tip, the URX dendrite makes a U-turn at the nose and loops back in the direction of the cell body in 74% or 65% of *hmn5* and *hmn6* adults, respectively, ultimately extending up to 150% of its normal length (Fig. 1B, Table I, and Fig. 2C). Further, the URX phenotype appears specific to dendrite length, as we did not observe defects in the expression of cell-specific markers, cell body positioning, dendrite branching, or other aspects of URX development. Finally, this phenotype is the opposite of the shortened dendrite phenotypes we had previously studied, and thus promised to shed light on a different aspect of dendrite growth control.

**Figure 2.**
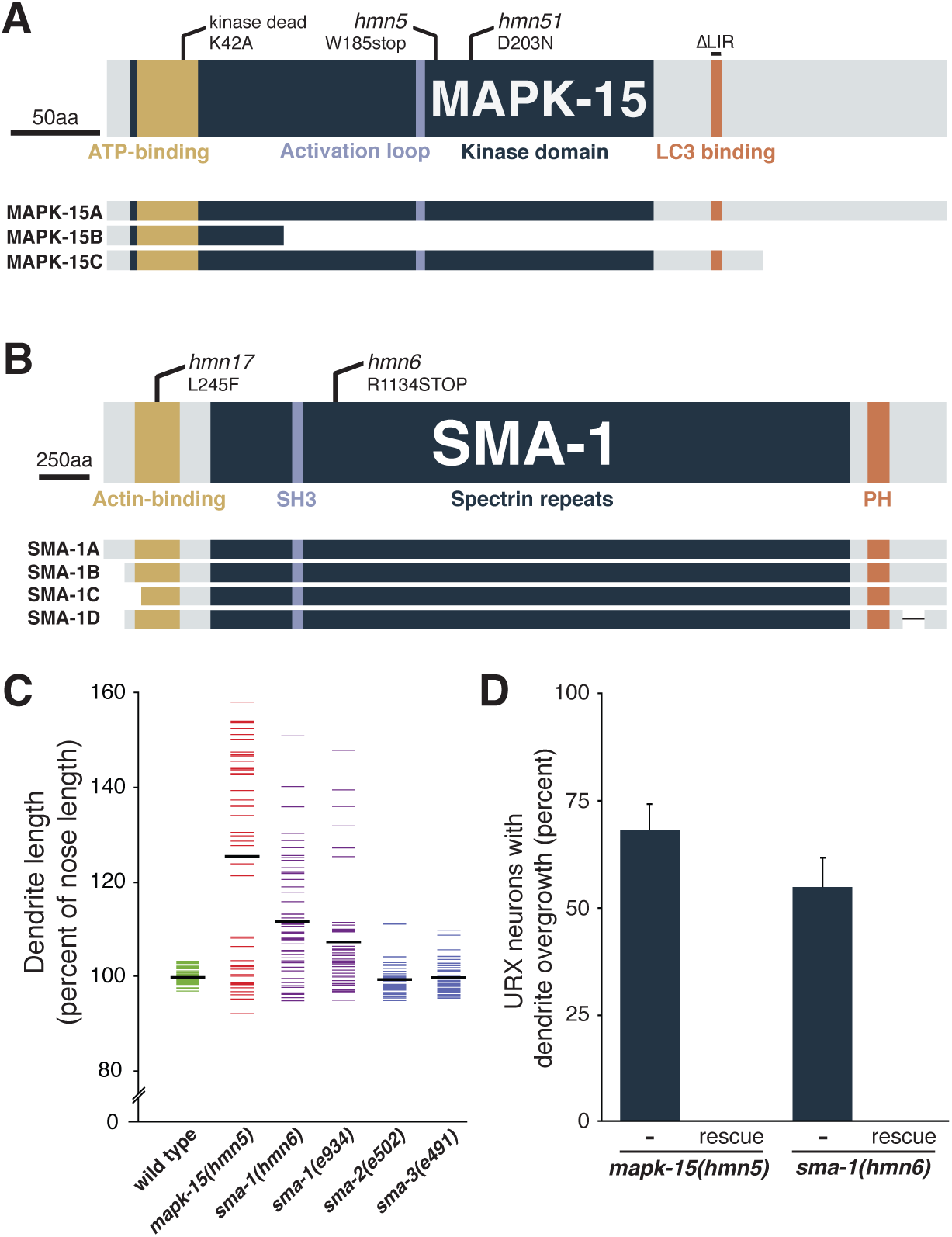
Loss of MAPK-15 or SMA-1 causes dendrite overgrowth. (A, B) Schematics of (A) the MAP kinase MAPK-15 and (B) the β_H_-spectrin SMA-1 showing effects of mutant alleles, conserved motifs and domains, and predicted isoforms. LC3 binding, conserved motif involved in binding autophagy protein LC3; SH3, Src Homology domain; PH, Pleckstrin Homology domain. (C) Animals expressing *flp-8*pro:GFP were synchronized at the fourth larval stage (L4) and dendrite and nose lengths measured. Each colored bar represents an individual; black bars represent population averages. n = 50 for each genotype, except n = 47 *sma-2(e502*). (D) Frequency of dendrite overgrowth was measured in *mapk-15* or *sma-1* mutants bearing the corresponding wild-type fosmids. Error bars, standard error of proportion. n = 50 for each genotype.

To identify the causative mutations in these strains, we performed whole-genome sequencing of pooled recombinants for *hmn5* and *hmn6*. *hmn5* contains a premature termination codon in the predicted gene C05D10.2 (Fig. 2A). A fosmid containing wild-type C05D10.2 completely rescues the phenotype (Fig. 2D). In a subsequent screen, we isolated another allele *hmn51* that also exhibits URX overgrowth (74% of animals, n=50; compare to Table I), fails to complement *hmn5*, and bears a point mutation in this gene (Fig. 2A). Together, these data indicate that disruption of C05D10.2 causes URX overgrowth. C05D10.2 is homologous to the mammalian MAP kinase ERK8/MAPK15, so we named this gene *mapk-15*. Recently, Piasecki et al., Kazatskaya et al., and Bermingham et al. independently identified roles for *mapk-15* in ciliated and dopaminergic sensory neurons of *C. elegans* [39–41]. As URX is neither ciliated nor dopaminergic, it is unclear how these roles relate to URX overgrowth.

*hmn6* contains a premature termination codon in *sma-1*, is completely rescued by a wild-type *sma-1* fosmid, and exhibits the characteristic small head and body size (Sma) phenotype associated with this gene (Fig. 2B,C) [42]. *hmn17* also exhibits the Sma phenotype, fails to complement *hmn6*, and contains a point mutation in *sma-1* (Fig. 2B). The previously described allele *sma-1(e934)* also exhibits URX overgrowth (Fig. 2C). *sma-1* encodes β_H_ spectrin, which is expressed in epithelial cells, links the cytoskeleton to the apical plasma membrane, and is required for normal embryonic elongation [43,44]. However, the *sma-1* dendrite defects are unlikely to be explained entirely by reduced head size, as *sma-2* and *sma-3* mutants also have reduced head size but do not exhibit comparable URX overgrowth (Fig. 2C; mean distance from URX cell body to nose (μm) ± SD: wild type, 112 ± 8; *sma-1(hmn6)*, 69 ± 6; *sma-1(e934)*, 78 ± 8; *sma-2(e502)*, 79 ± 5; *sma-3(e491)*, 86 ± 5).

### URX dendrite overgrowth increases with age

To understand when *mapk-15* and *sma-1* act, we examined the extent of URX overgrowth at different developmental stages. We very rarely observed dendrite overgrowth in first larval stage (L1) animals, suggesting overgrowth is not primarily due to defects in initially shaping the dendrite during embryonic development (Fig. 3A,F). Rather, dendrite overgrowth increases with age, reaching moderate penetrance in L4 animals and becoming even more pronounced in two-day adults (penetrance in L1, L4, 2A: 4%, 44%, 74% in *mapk-15(hmn5)*; 8%, 17%, 65% in *sma-1(hmn6);* Fig. 3). The extent of the overgrowth also increases with age (Fig. 3F). Moreover, rare adult *mapk-15* animals on crowded, starved plates exhibit profusely overgrown dendrites that include ectopic branching, a phenotype we never observed in wild-type animals (Fig. 3D). These results suggest that *mapk-15* and *sma-1* act post-embryonically to constrain dendrite growth throughout the life of the animal.

**Figure 3.**
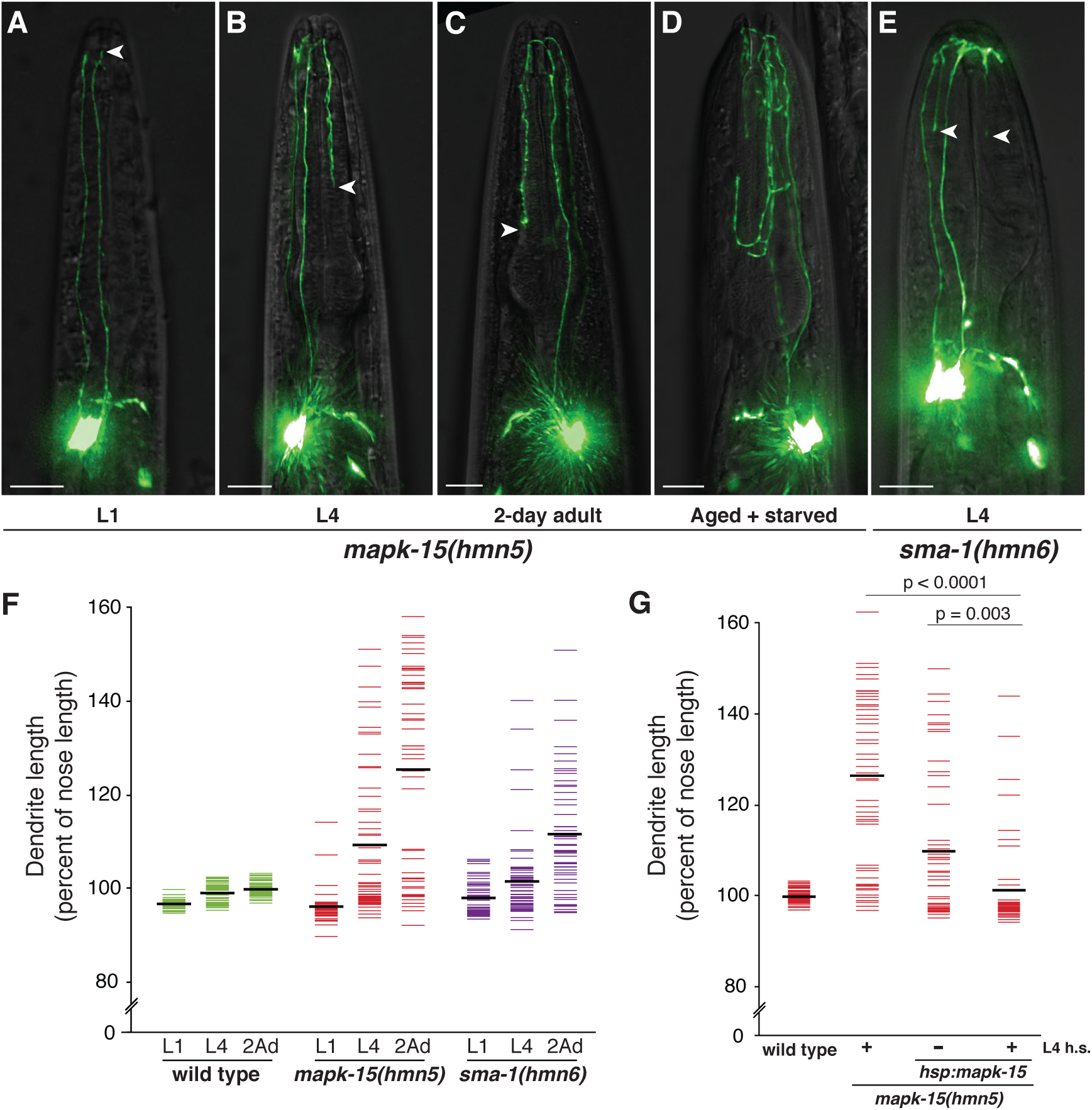
Dendrite overgrowth is age-dependent. (A-E) *mapk-15* or *sma-1* mutants at the indicated developmental stages, or from ~2-week old starved plates, were imaged. Arrowheads, dendrite endings. Scale bars, 10 μm. (F) Wild-type, *mapk-15*, or *sma-1* animals were synchronized at the L1 or L4 stages or as two-day adults (2Ad) and dendrite lengths and nose lengths were measured. (G) *mapk-15* mutants bearing a transgene encoding a heat-shock-inducible *mapk-15(+) (hsp:mapk-15)* were synchronized at the L4 stage, subjected to a brief heat shock (+ L4 h.s., 30 min, 34°C) or not (– L4 h.s.), recovered, and dendrite and nose lengths were measured in two-day adults. p-values, Mann-Whitney U-test. Colored bars are individual animals, black bars are population averages. n ≥ 50 in all cases.

To test this hypothesis, we generated constructs in which a wild-type *mapk-15* genomic fragment is placed under control of heat-shock-inducible promoters (*hsp*, see Methods). We did not perform this experiment with *sma-1* due to its larger gene size (~12 kb). We grew *mapk-15(hmn5)* animals bearing these transgenes, induced expression of wild-type *mapk-15* in L4 animals, and then measured URX dendrite length in two-day adults. We observed a strong, albeit incomplete, rescue of the overgrowth defect (Fig. 3G). These results indicate that *mapk-15* is not necessary during early URX development to prevent overgrowth, but its expression at the time when the phenotype appears is sufficient to restrict growth. Induction of expression in embryos or two-day adults showed a modest (p=0.08) or not appreciable (p=0.94) effect, respectively (Supp. Fig. S2). These results suggest that *mapk-15* may temporally regulate URX dendrite growth.

### *mapk-15* acts cell-autonomously to constrain dendrite growth

Next, we wanted to know in which cells *mapk-15* and *sma-1* act to regulate dendrite growth. Previous studies had shown that *sma-1* is expressed primarily in hypodermis, intestine, and pharynx [43,44]. To determine where *mapk-15* is expressed, we generated a transcriptional reporter and found that it labels URX as well as many other head sensory neurons at all larval stages (Fig. 4A; expression in ciliated sensory neurons is also reported in [39–41]), suggesting *mapk-15* could act cell-autonomously in URX.

**Figure 4.**
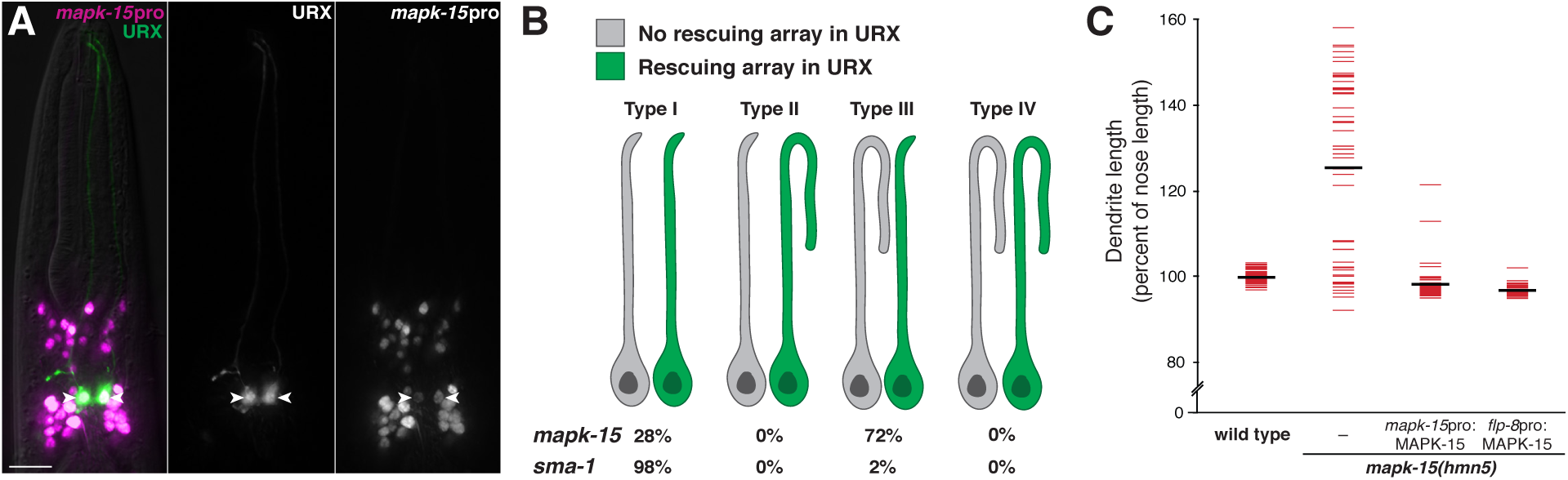
*mapk-15* acts cell-autonomously in URX to constrain dendrite growth. (A) Image of a wild-type animal expressing *flp-8*pro:GFP (URX, green) and *mapk-15*pro:NLS-mCherry (magenta). Arrowheads, URX cell bodies. Scale bar, 10 μm. (B) Mosaic analysis was performed on *mapk-15* or *sma-1* mutants with integrated *flp-8*pro:GFP (URX) and bearing unstable extrachromosomal arrays with *flp-8*pro:mCherry and wild-type *mapk-15(+)* or *sma-1(+)* respectively. Presence of mCherry in only the left or right URX was used to identify genetic mosaics, and the dendrite overgrowth was scored in both the neuron carrying (green) and lacking (gray) the array. Proportion of animals in each class is shown. (C) Wild-type or *mapk-15* animals bearing transgenes consisting of wild-type *mapk-15* under control of its own promoter (*mapk-15*pro:MAPK-15), a URX-specific promoter (*flp-8*pro:MAPK-15), or neither (–) were synchronized as two-day adults and dendrite and nose lengths were measured. Colored bars, individual dendrites; black bars, population averages. n ≥ 44 in all cases.

To determine if *mapk-15* and *sma-1* normally act in URX, we performed mosaic experiments in which mutant animals carried an extrachromosomal array bearing a fosmid encompassing the wild-type gene together with a fluorescent marker that allowed us to score the presence or absence of the transgene in URX. Such extrachromosomal arrays are stochastically lost during cell division, which creates genetic mosaics. We selected adult mosaic animals in which the array was present in one URX neuron but absent in the contralateral URX neuron, and assessed dendrite overgrowth. We considered four possible outcomes: (I) overgrowth in <u>neither neuron</u>; (II) overgrowth only in the neuron <u>with the array</u>; (III) overgrowth only in the neuron <u>without the array</u>; or (IV) overgrowth in <u>both neurons</u> (Fig. 4B).

For *mapk-15* mosaics, we found that 72% of animals were type III and 28% of animals were type I (Fig. 4B). Thus, when the rescuing array is present in URX, the dendrite is always normal length, and when the array is absent from URX, the dendrite exhibits overgrowth at the previously observed penetrance of the *mapk-15* phenotype (compare with 74%, Table I). This result suggests that *mapk-15* acts cell-autonomously in URX to constrain dendrite growth. In contrast, for *sma-1* mosaics, we found that 98% of animals were type I, suggesting that the overgrowth phenotype was efficiently rescued regardless of the presence or absence of the array in URX (Fig. 4B), and thus *sma-1* likely acts in other cells to constrain URX dendrite growth.

An advantage of mosaic experiments is that the rescuing transgene is expressed under control of its native regulatory sequences; however, because these experiments rely on loss of the transgene during cell division, they can fail to resolve cells that are closely related by developmental lineage. Therefore, to further test whether *mapk-15* can function cell-autonomously in URX, we expressed a transgene containing a wild-type *mapk-15* genomic fragment under control of its own promoter or a URX-specific promoter (*flp-8*pro) in mutant animals and measured dendrite lengths in two-day adults. We found that expression of *mapk-15* with either promoter rescued the overgrowth phenotype (Fig. 4C). Together, these results suggest that *sma-1* likely acts non-cell-autonomously from neighboring tissues, whereas *mapk-15* is normally expressed in URX and its expression in URX is necessary and sufficient to constrain dendrite growth.

### MAPK-15 localizes to the URX dendrite ending and requires its kinase activity

To gain insight into how MAPK-15 regulates URX dendrite growth, we asked where it localizes in the neuron and whether it requires its predicted kinase activity. We generated a construct encoding a superfolderGFP(sfGFP)-MAPK-15 fusion protein (see Methods) and expressed it in URX. This construct rescues the *mapk-15* dendrite phenotype, indicating the fusion protein is functional (Supp. Fig. S3A). We found that sfGFP-MAPK-15 localizes throughout the neuron with ~4-fold relative enrichment at the URX dendrite ending (relative fold enrichment ± SD, 3.7 ± 2.0), suggesting it may act locally to constrain growth (Fig. 5A-B, Supp. Fig. S3C-D). Notably, many neuronal receptive endings – including cilia of *C. elegans* sensory neurons and dendritic spines in mammals – are known to contain signaling factors that regulate their own growth, typically on the scale of 1-5 μm [3,45]. In these examples, the receptive ending is a discrete compartment that is biochemically isolated from the rest of the dendrite. In contrast, the URX dendrite is not ciliated and ultrastructural studies have not revealed any obvious physical compartmentalization at its ending. However, guanylate cyclase proteins that mediate URX sensory functions (e.g. GCY-35) have been shown to localize to the dendrite ending, raising the possibility that the dendrite contains a subcellular compartment at its ending that is specialized for sensory signaling [33].

**Figure 5.**
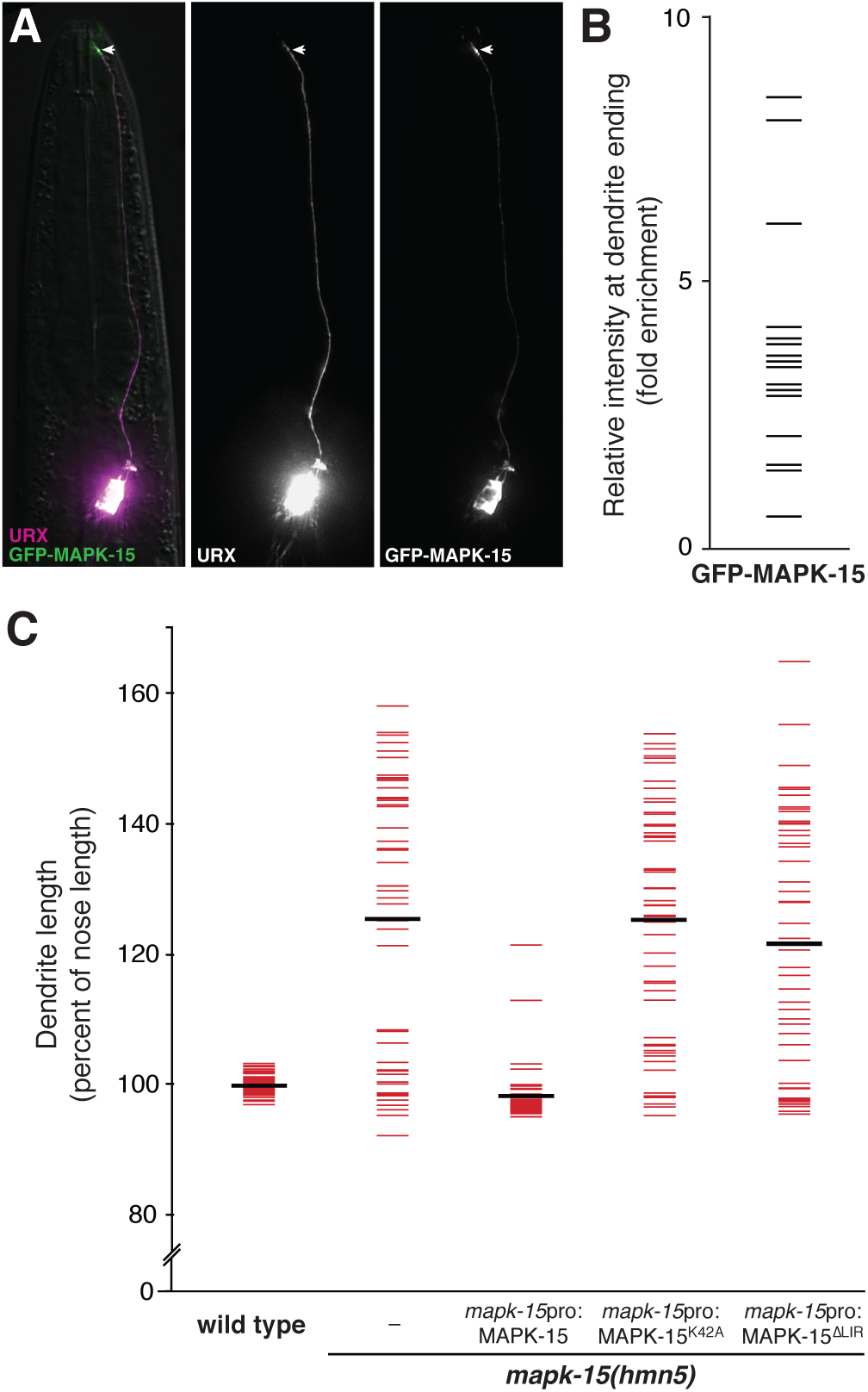
MAPK-15 localizes to dendrite endings and requires its kinase activity and LIR motif. (A) Wild-type L4 animal expressing *flp-8*pro:mCherry (URX) and *flp-8*pro:superfolderGFP-MAPK-15 (URX). Arrow, dendrite ending. (B) Relative enrichment of superfolderGFP-MAPK-15 at the dendrite ending compared to its middle, as shown in Supp. Fig. S4. Mean enrichment ± SD, 3.7 ± 2.0. n = 20. (C) Wild-type or *mapk-15* animals bearing no transgene (–) or a transgene encoding wild-type, kinase-dead (K42A), or ∆LIR alleles of *mapk-15* under its own promoter were synchronized as two-day adults and dendrite and nose lengths were measured. Colored bars, individual animals; black bars, population averages. n = 50 in all cases.

To test whether MAPK-15 might participate in localized signaling, we asked whether it functions as a kinase. MAPK-15 has several conserved domains that are typical of MAP kinases, including an ATP-binding pocket (Fig. 2B). Previous work on mammalian MAPK15 has shown that a conserved lysine residue in this domain is necessary for kinase activity [46,47]. We therefore generated a presumed kinase-dead allele (MAPK-15(K42A), see Methods) under the control of its own promoter, introduced it as a transgene into mutant animals, and asked whether it could rescue dendrite overgrowth. We observed no rescue, consistent with the idea that MAPK-15 requires its kinase activity (Fig. 5C). We also generated an allele lacking the LC3-interacting region (MAPK-15(∆LIR)), a motif that has been shown in mammalian MAPK15 to bind factors involved in autophagy [48,49]. This allele also failed to rescue (Fig. 5C). While we cannot exclude kinase- or LIR-dependent roles for MAPK-15 elsewhere in the cell, taken together these results suggest that MAPK-15 may constrain dendrite growth through localized signaling or regulatory interactions at the dendrite ending.

### The guanylate cyclase GCY-35 localizes throughout the expanded dendrite ending

Given that GCY-35 normally localizes to the URX dendrite ending near the nose, we wondered where it would localize in an overgrown dendrite. We considered three possibilities: it might localize to the overgrown dendrite ending; to the region of the dendrite closest to the nose; or, throughout the overgrown portion of the dendrite. The first two possibilities would suggest that *mapk-15* regulates the growth of the dendrite but not of the sensory region, whereas the third possibility suggests an expansion of the sensory region itself.

Therefore, to distinguish these possibilities, we examined the localization of GCY-35-GFP in URX in wild-type and *mapk-15* animals. We found that GCY-35-GFP was enriched at the dendrite ending in wild-type animals, consistent with previous reports, as well as in *mapk-15* young animals prior to the appearance of the dendritic overgrowth (Fig. 6A). However, in *mapk-15* L4 animals in which the dendritic overgrowth was apparent, GCY-35-GFP localized throughout the overgrown region (Fig. 6B). Its localization also expanded into more proximal regions of the dendrite where it is not normally enriched (Fig. 6B). We obtained similar results with the globin GLB-5, which regulates oxygen sensing and also localizes to the wild-type URX dendrite ending (Supp. Fig. S4A) [33]. These results are consistent with the notion that dendritic overgrowth corresponds to expansion of a subcellular sensory region at the dendrite ending.

**Figure 6.**
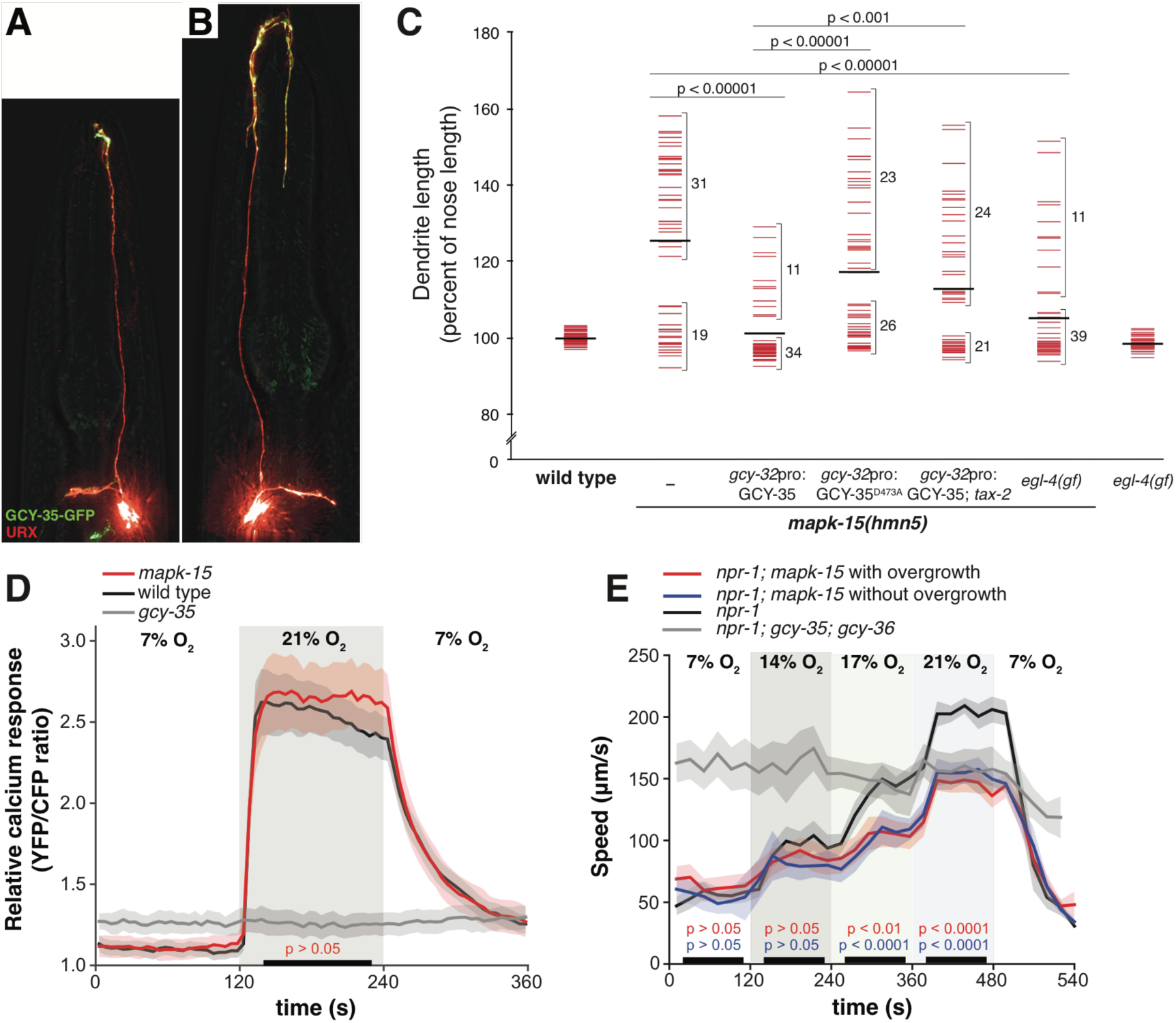
Dendrite overgrowth may reflect expansion of a sensory compartment. (A, B) *mapk-15* mutants expressing the oxygen-sensing guanylate cyclase GCY-35-HA-GFP and soluble mCherry under control of a URX-specific promoter (*gcy-37*pro) were imaged (A) at the L3 stage before overgrowth or (B) at the L4 stage after overgrowth. (C) Wild-type, *mapk-15*, *mapk-15; tax-2*, *egl-4(gf),* and *mapk-15;egl-4(gf)* animals were synchronized as two-day adults and dendrite and nose lengths were measured. *mapk-15* and *mapk-15; tax-2* mutants carried transgenes overexpressing wild-type GCY-35 or a cyclase-dead mutant (GCY-35^D473A^). Colored bars, individual dendrites; black bars, population averages. Brackets indicate numbers of dendrites in the indicated range. (D) URX calcium responses to a 7% – 21% oxygen shift were measured in wild-type (n = 18), *mapk-15* animals with dendrite overgrowth (n = 17), and *gcy-35* animals (n = 10) as sensory-defective controls. Solid lines, average YFP/CFP ratio; shaded areas, SEM; black bars, time intervals used for statistical comparison vs wild-type control. n=15 in all cases. (E) Locomotory activity was measured in response to 7% – 14% – 17% – 21% oxygen shifts for *npr-1* animals, *npr-1; mapk-15* animals with or without URX overgrowth, and *npr-1; gcy-35; gcy-36* animals as sensory-defective controls. Solid lines, average speed; shaded areas, SEM; black bars indicate time intervals used for statistical comparisons. Color-coded p-values, *mapk-15* with/without overgrowth vs *npr-1* controls (n > 70). Comparison of *mapk-15* animals with overgrowth vs those without overgrowth yielded p>0.05 for all conditions. In all panels, p-values were calculated with the Mann-Whitney U test.

### URX dendrite overgrowth is suppressed by genetically mimicking increased cGMP signaling

While examining GCY-35-GFP localization, we were surprised to find that overexpression of this transgene suppressed the *mapk-15* overgrowth phenotype (Fig. 6C, penetrance with and without GCY-35-GFP overexpression, 62% and 24% respectively, p<0.0001). This result raises the possibility that elevated guanylate cyclase function might mitigate the effects of *mapk-15* disruption.

We were concerned that overexpression of GCY-35 might cause phenotypic changes for reasons unrelated to its guanylate cyclase activity. Therefore, we performed two additional controls. First, we introduced a point mutation in GFP-GCY-35 that inactivates its catalytic site [31]. We found that overexpression of this transgene in URX failed to suppress the *mapk-15* overgrowth (Fig. 6C), indicating that suppression requires the guanylate cyclase activity. Second, we reasoned that overexpression of an active guanylate cyclase is likely to increase cellular cGMP levels, which could then act on cGMP-dependent protein kinases and cyclic nucleotide-gated (CNG) channels [50]. Therefore, we asked whether the CNG channel TAX-2 is important for GCY-35-mediated suppression of dendrite overgrowth. Indeed, we found that in *mapk-15; tax-2* mutant animals, GCY-35-mediated suppression of overgrowth was partially relieved (Fig. 6C).

Finally, we wanted to use an independent genetic manipulation to mimic elevated cGMP levels and ask whether that would also suppress dendrite overgrowth. For this purpose, we took advantage of a gain-of-function mutation in the cGMP-dependent protein kinase EGL-4 [*egl-4(gf)*] [51]. Consistent with the results above, *egl-4(gf)* efficiently suppressed the *mapk-15* dendrite overgrowth (Fig. 6C, penetrance with and without *egl-4(gf)*, 62% and 22% respectively, p<0.00001). This effect is specific to *egl-4* hyperactivity, as an *egl-4* loss-of-function mutation did not alter URX dendrite length or enhance the *mapk-15* overgrowth (Supp. Fig. 2). Together, these results suggest that genetically mimicking increased cGMP signaling suppresses dendrite overgrowth.

### URX dendrite overgrowth is not correlated with overt defects in oxygen sensation

Sensory cilia can increase in length when deprived of sensory input, as if trying to compensate for reduced sensory signaling [4]. Although URX is not ciliated, we wondered whether *mapk-15* mutants might have reduced sensory signaling relevant to dendrite overgrowth. Rising oxygen levels evoke a tonic increase in intracellular calcium levels in URX neurons [35,52]. We introduced a genetically-encoded calcium reporter, yellow cameleon 2.60, into *mapk-15* animals and used 2-day adults exhibiting dendrite overgrowth to visualize calcium changes in the URX cell body in response to a 7% - 21% oxygen shift. As a control, sensory-defective *gcy-35* mutant animals completely lack an oxygen response (Fig. 6D). In contrast, we found that oxygen-evoked calcium responses in the *mapk-15* mutant were indistinguishable from those of wild-type animals (Fig. 6D).

We also measured behavioral responses to changes in ambient oxygen levels. Because the standard laboratory reference strain (N2) carries a gain-of-function mutation in the neuropeptide receptor NPR-1 that suppresses behavioral responses to oxygen, we performed our behavioral assays in strains carrying an *npr-1* loss-of-function mutation that mimics the behavior of true wild isolates [32, 53–55]. These *npr-1* animals increase their speed of locomotion and re-orient their direction when challenged with elevated oxygen concentrations [32,53]. This behavior depends on URX and the oxygen sensors GCY-35 and GCY-36 [32,53]. We found that *mapk-15* mutants have overall slower speed in response to elevated oxygen, but these responses are equivalent among mutant animals whether or not they exhibit the URX dendrite overgrowth. Thus, this altered behavioral change is unlikely to be caused by URX dendrite length and probably reflects other disruptions to organismal sensory function or general physiology in the mutant.

While these experiments cannot exclude subtle alterations in URX sensory processing, or the possibility that larger sensory defects are masked by adaptation in the neuron, taken together these data do not support the idea that dendrite overgrowth is a response to URX sensory deficits. Conversely, other mutants that cause URX sensory deficits have not been reported to lead to overgrowth, and we did not observe overgrowth with *egl-4(lf)* (Supp. Fig. S4B). Therefore, our results suggest that overgrowth is unlikely to be a secondary consequence of URX sensory deficits, and thus point to a more direct role for MAPK-15 in regulating dendrite length.

## DISCUSSION

### MAPK-15 and the control of sensory dendrite length

How cells measure their own size is a basic problem in cell biology [56]. Here, we show that MAPK-15 acts in the URX neuron to constrain the length of its sensory dendrite. In the absence of MAPK-15, the dendrite exhibits age-dependent overgrowth. MAPK-15 acts temporally when the overgrowth first appears and spatially at the dendrite ending. These results suggest that MAPK-15 acts locally to constrain dendrite length.

Two lines of evidence suggest that dendrite overgrowth corresponds to expansion of a subcellular sensory compartment. First, the oxygen-sensing guanylate cyclase GCY-35, which normally marks a subcellular region at the dendrite ending, expands its localization to fill the overgrowth. Second, genetic manipulations that mimic hyperactive sensory signaling can suppress the overgrowth. Intriguingly, three recent studies described a role for MAPK-15 in other information-processing subcellular compartments in neurons. Piasecki et al., Kazastaya et al., and Bermingham et al. report localization of MAPK-15 in sensory cilia and at dopaminergic synapses of certain ciliated sensory neurons [39–41]. Loss of MAPK-15 leads to altered sensory cilia morphology, defects in mating behaviors mediated by ciliated sensory neurons and defects in swimming behaviors affected by dopamine signaling [39–41]. While URX is not ciliated, these results are consistent with the idea that MAPK-15 acts at subcellular sensory compartments.

How might MAPK-15 control dendrite length? We can imagine two general models. First, the dendrite ending might be a relatively static structure, and loss of MAPK-15 creates an aberrant positive signal that tells the neuron to “make more dendrite.” Alternatively, the dendrite ending might be in a state of dynamic equilibrium, where its steady-state length reflects a balance between the addition and retrieval of material. In this model, loss of MAPK-15 might shift the balance towards addition, leading to overgrowth, whereas genetically mimicking hyperactive sensory signaling might reduce the rate of addition, shifting the balance back. Our current data cannot distinguish between these models. However, it is interesting to note some circumstantial evidence that may point to the idea that MAPK-15 normally promotes retrieval of dendrite material. First, MAPK15/ERK8 plays a role in autophagy in other systems [48,49]. Second, its interaction with autophagic components is mediated through its conserved LC3-interacting region [49], which we found to be required to restrict dendrite growth. Third, in *C. elegans* ciliated neurons, MAPK-15 localizes near the periciliary membrane compartment, a region that is enriched for proteins involved in membrane trafficking [40,41]. Fourth, the role of MAPK-15 in *C. elegans* dopaminergic neurons has been suggested to involve membrane trafficking of a dopamine transporter at synapses [39]. Perhaps an altered balance of membrane delivery and retrieval underlies the defects at the URX dendrite ending as well as the phenotypes observed in other cell types.

### Age-dependent dendrite remodeling

Our results define a genetic pathway that affects dendrite length throughout the life of the organism. Little is known about how dendrite arbors remodel with age in mammalian brain. Many studies have focused on changes in dendritic spines, while relatively few have examined changes in dendrite length or branching [57]. Classical studies using Golgi staining to examine dendrite arbors in mammalian cortex reported a reduction in dendrite length and branching with age, including in humans [58,59]. The cause of this change – and whether it is simply "deterioration" – is not known. Paradoxically, some studies have reported dendrite branching to increase rather than decrease with age [60,61]. An intriguing possibility is that age-dependent dendritic changes reflect the overall balance of dendrite growth and retraction in mature life, which may differ between brain regions or neuron types. Indeed, time-lapse imaging of cortical neurons showed that some dendrite arbors remain highly dynamic during adulthood [62]. The mechanisms that control dendrite length throughout life have remained unclear.

In *C. elegans*, a few examples of post-embryonic dendrite remodeling have been demonstrated. During the dauer larval stage, the unbranched dendrites of IL sensory neurons are converted into highly branched dendritic arbors [63]. The mechanosensory neurons PVD and FLP elaborate a branched dendritic arbor beginning in young larvae that becomes increasingly complex as development progresses [20]. Age-dependent changes in dendrite arborization have been carefully examined in PVD [64]. In older adults, the PVD arbor becomes increasingly hyper-branched and disorganized, reminiscent of some of the age-dependent changes in URX that we observe in *mapk-15* mutants [64]. Throughout development, the PVD dendrite arbor is highly dynamic, and its branching pattern reflects the net result of many outgrowth and retraction events [17,20,64]. Interestingly, in both younger and older animals (L4 larvae and 5-day adults), PVD dendrite outgrowth is slightly favored over retraction [64]. The accumulated effects of this imbalance may explain its age-dependent hyper-branching. This example would be consistent with a dynamic equilibrium model for control of dendrite length in URX and, possibly, in other systems.

Dendrite morphogenesis has been studied in *C. elegans* in the context of branching (PVD, FLP, and IL neurons) and DYF-7-mediated retrograde extension (ciliated sensory neurons of the amphid) [12– 23,30,63]. Here, we show that URX offers a complementary model of dendrite morphogenesis that provides new mechanistic insights. In particular, our results show that, in addition to established pathways that promote dendrite growth and branching, there are also yet-unidentified pathways that constrain excessive dendrite growth throughout the life of the animal. It will be important to determine whether these age-dependent changes reflect a shift in the rates of addition and removal of dendrite material, and how they relate to alterations in dendrite lengths observed in the aging mammalian brain.

## METHODS

### DNA constructs and transgenes

The following DNA constructs and transgenes were used: *ynIs78*[*flp-8*pro:GFP] X and *flp-8*pro [36]; *hsp-16*.*41*pro and *hsp-16.2*pro (from pPD49.83 and pPD49.78, Andrew Fire); *mapk-15*pro (1.7 kb upstream of coding sequence); *gcy-32*pro [65]; *mapk-15* (genomic fragment); MAPK-15(K42A) (codon change generated via overlap extension PCR from *mapk-15* genomic fragment); MAPK-15(∆LIR) (deletion of codons 342-345 (encoding YEMI) generated via PCR from *mapk-15* genomic fragment); *gcy-32*pro, *gcy-37*pro, GCY-35-GFP, GCY-35-HA-GFP [33]; GCY-35(D473A)-GFP (codon change generated via overlap extension PCR from GCY-35-GFP fragment), superfolderGFP [66]; *gcy-37*pro and yellow cameleon YC2.60 [33,67]. SuperfolderGFP is a robustly folding GFP variant that was used for convenience [66]. The *mapk-15* genomic region likely contains some transcriptional regulatory sequences, as the *flp-8*pro:superfolderGFP-*mapk-15* construct shows weak expression in an unidentified head neuron not seen with *flp-8*pro:GFP alone; however, expression in this cell does not interfere with imaging and does not appear to be sufficient for rescue based on mosaic analysis and the phenotype observed in strains bearing *hsp:mapk-15* in the absence of heat shock. Strains are listed in Supp. Table I.

### Genetic screen and mutant identification

L4 stage animals were mutagenized using 70 mM ethyl methanesulfonate (EMS, Sigma) at approximately 22°C for 4 hours [42]. Nonclonal F2 progeny were examined on a Nikon SMZ1500 stereomicroscope with an HR Plan Apo 1.6x objective, and animals with aberrant dendrite morphologies were recovered to individual plates. Mutant identification was performed by one-step pooled linkage analysis [68] and sequence variants were analyzed with Cloudmap [69]. Identified mutations are listed in Supp. Table II.

### Heat shock

Animals bearing *hsp*:*mapk-15* constructs (Supp. Table I) were subjected to heat shock by incubating them at 34°C for 30 min on standard growth medium in the presence of bacterial food. Even in the absence of heat shock, these constructs mildly rescued dendrite defects, possibly due to "leaky" expression (compare Fig. 3F (2Ad) with Fig. 3G (-h.s.)).

### Microscopy and image processing

Animals were mounted on agar pads and immobilized with sodium azide. Image stacks were collected on a DeltaVision Core deconvolution imaging system (Applied Precision) with an InsightSSI light source; a UApo 40x/1.35 NA oil immersion objective, PlanApo 60x/1.42 NA oil immersion objective, or UPlanSApo 100x/1.40 NA oil immersion objective; the Live Cell Filter module; and a Photometrics CoolSnap HQ2 CCD camera (Roper Scientific). Image stacks were acquired and deconvolved with Softworx 5.5 (Applied Precision). Maximum projections were generated with contiguous optical sections in ImageJ (NIH), then linearly adjusted for brightness in Adobe Photoshop. Multicolored images were created by placing each channel in a separate false-colored screen layer in Photoshop. Figures were assembled in Adobe Illustrator.

### Quantification of dendrite lengths and statistical tests

Dendrite lengths were measured using the segmented line tool in ImageJ (NIH). The dendrite was traced from the point where it joins the cell body to the point where it ends at the nose, then normalized by the distance from the cell body to the nose to account for variance in the size of the animal. The data were initially recorded in Microsoft Excel or Apple Numbers, and statistical analysis was performed using R version 3.4.2 (https://www.r-project.org/). To test statistical significance, the Mann-Whitney U test was chosen to compare differences between independent samples that are not necessarily normally distributed.

### Calcium imaging

A *gcy-37*pro*:YC2.60* (yellow cameleon 2.60) transgene was used for ratiometric imaging of relative calcium concentration in URX cell bodies [33,67]. L4 animals expressing the sensor were picked 48h before imaging. Calcium imaging experiments were performed as described previously [33,67] using an inverted microscope (Axiovert, Zeiss) equipped with a 40x C-Apochromat lens (Zeiss), Optosplit II beam splitter (Optical Insights), Evolve Delta camera (Photometrics) and MetaMorph acquisition software (Molecular Devices). Photobleaching was minimized using a 2.0 optical density filter. Animals were glued to agarose pads (2% agarose in M9 buffer, 1 mM CaCl_2_) using Dermabond tissue adhesive (Ethicon) with the nose immersed in a mix of bacterial food (*E. coli* OP50) and M9 buffer. To deliver gas stimuli, glued animals were placed under a Y-shaped microfluidic chamber with inlets connected to a PHD 2000 Infusion syringe pump (Harvard Apparatus) running at a flow rate of 2.5 ml/min. An electronic valve system placed between the syringes and the microfluidic chamber allowed switching between two different gas mixtures in a controlled manner at pre-specified time intervals. URX calcium responses were recorded at 2 frames/s with an exposure time of 100 ms. Image and statistical analysis was performed using Neuron Analyser, a custom-written MATLAB program (code available at https://github.com/neuronanalyser/neuronanalyser). Statistical comparisons of calcium responses were done using a Mann–Whitney U test.

### Behavioral assays

Locomotion was assayed as described previously [34], with slight alterations. L4 animals were picked 48 h before the assay. Two days later, 20-30 adult hermaphrodites were transferred to nematode growth medium plates containing low peptone (5% of standard bactopeptone concentration) that had been seeded 16–20 h earlier with 20 μl of *E. coli* OP50 grown in 2× TY medium (per liter, 16 g tryptone, 10 g yeast extract, 5 g NaCl, pH 7.4). To deliver gas stimuli, animals were placed under a 1 cm × 1 cm × 200 μm deep polydimethylsiloxane chamber with inlets connected to a PHD 2000 Infusion syringe pump (Harvard apparatus). Humidified gas mixtures were delivered at a flow rate of 3.0 ml/min. We recorded movies using FlyCapture (Point Gray) on a Leica M165FC dissecting microscope with a Point Gray Grasshopper camera running at 2 frames/s. Movies were analyzed using Zentracker, a custom-written MATLAB software (code available at https://github.com/wormtracker/zentracker). Speed was calculated as instantaneous centroid displacement between successive frames. Locomotion assays were done in triplicate on at least two independent days. For statistical comparisons, we chose time intervals where we expected the speed changes to have plateaued, that is, with a delay with respect to the timing of the switch in O_2_ concentration. For the intervals of interest, we determined independent per-subject means derived from individuals flagged as continuously valid for at least 10 s during the interval. We considered all individuals in the field of view as valid except those in contact with other animals and those that were off the food lawn or less than half a body-length from the border. Following these criteria, each individual was sampled at most once per interval; n indicates the minimum number of valid samples obtained per interval. Statistical comparisons for speed assays used a Mann-Whitney U test.

## ACKNOWLEDGMENTS

We thank the CGC, which is funded by NIH Office of Research Infrastructure Programs (P40 OD010440), and WormBase.

---

**Supplemental Figure S1.**
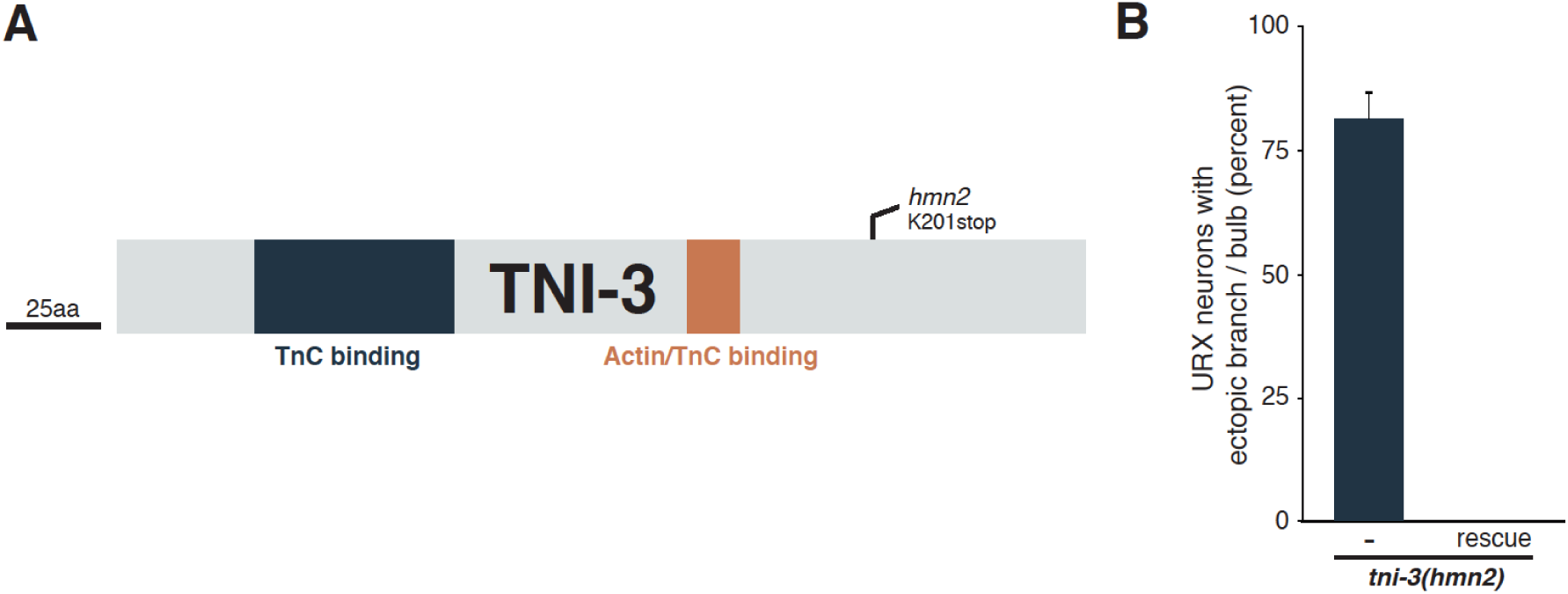
Loss of TNI-3 causes ectopic branch formation. (A) Schematic of the Troponin I TNI-3 showing effects of the mutant allele *hmn2* and conserved motifs and domains. TnC, Troponin C. (B) Frequency of dendrite overgrowth was measured in *tni-3* mutants bearing the wild-type fosmid. Error bars, standard error of proportion. n ≥ 15.

**Supplemental Figure S2.**
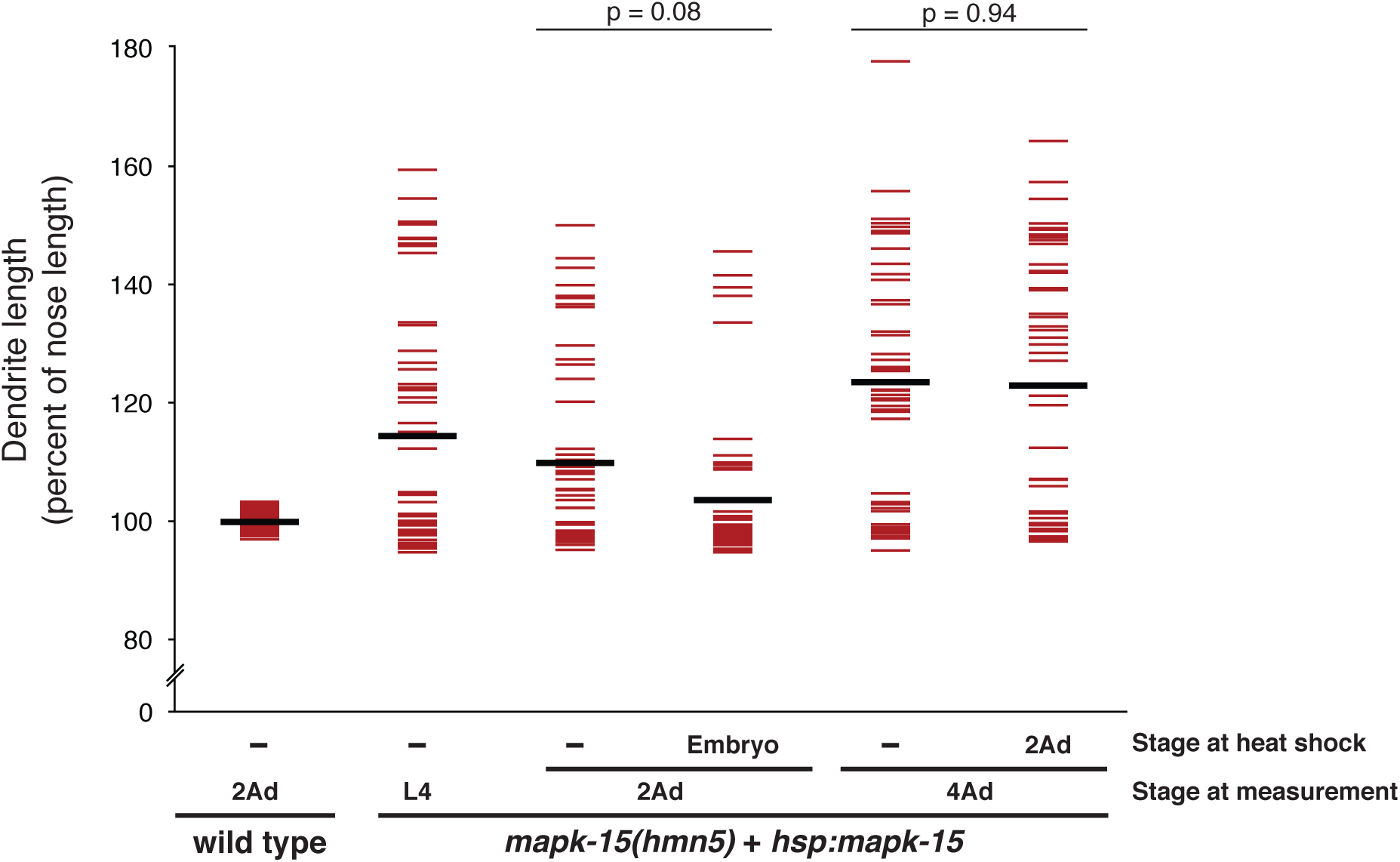
Induction of MAPK-15 expression at early or late developmental timepoints does not efficiently rescue dendrite overgrowth. *mapk-15* mutants bearing a transgene encoding a heat-shock-inducible *mapk-15(+)* genomic fragment *(hsp:mapk-15)* were subjected at the indicated developmental stage to a brief heat shock (30 min, 34°C) or not (–), recovered, and dendrite and nose lengths were measured at the indicated stage. Embryo, mixed-stage embryos. 2Ad, second day of adulthood. 4Ad, fourth day of adulthood. p-values, Mann-Whitney U-test. Colored bars are individual animals, black bars are population averages. n ≥ 50 in all cases.

**Supplemental Figure 3.**
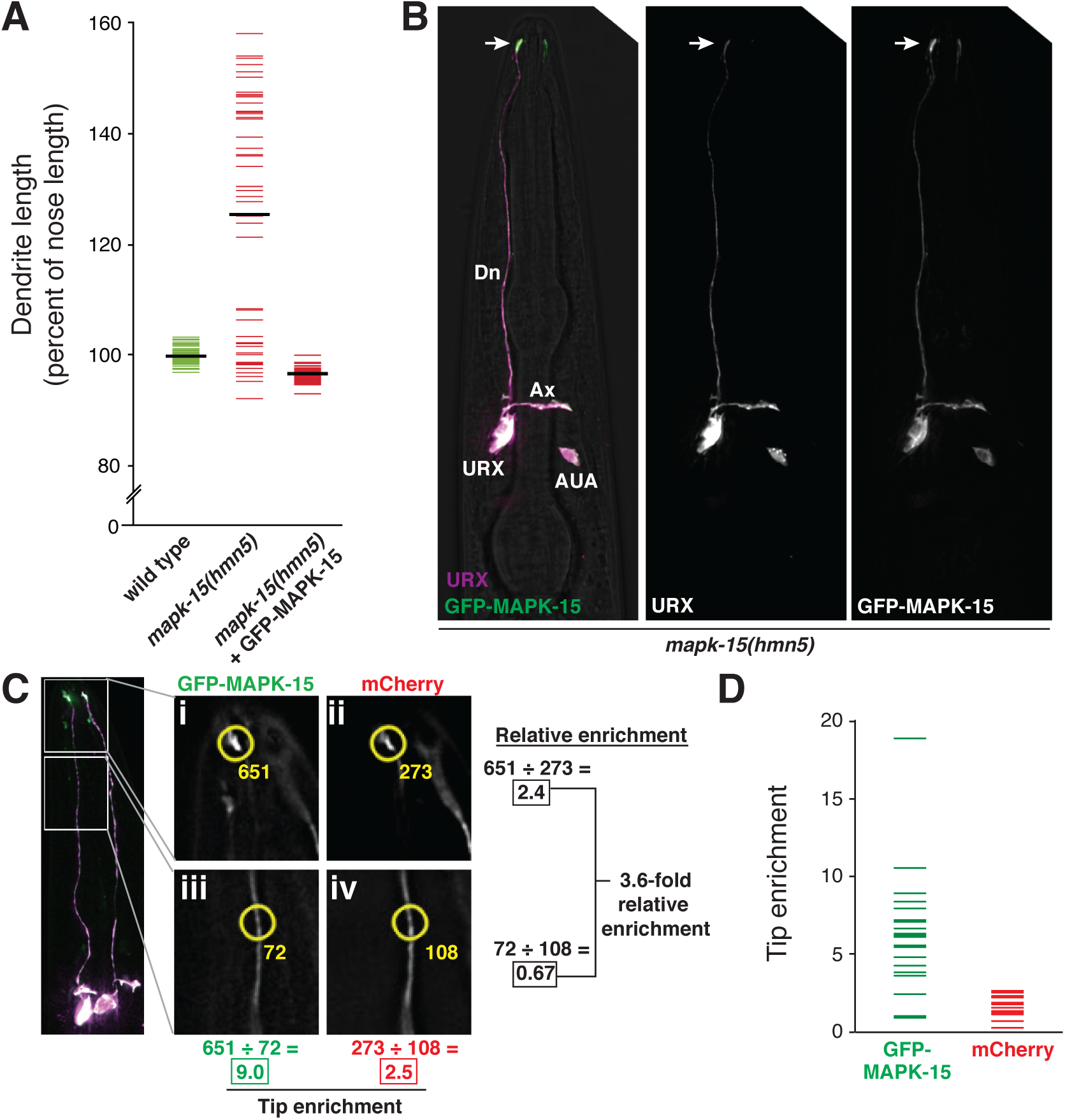
A functional GFP-MAPK-15 is enriched at the dendrite ending. (A) Wild-type or *mapk-15* animals with or without *flp-8*pro:superfolderGFP-MAPK-15 were synchronized as two-day adults and dendrite and nose lengths were measured. Colored bars, individual animals; black bars, population averages. n = 50 in all cases. (B) *mapk-15* mutant animal expressing *flp-8*pro:mCherry (URX) and *flp-8*pro:superfolderGFP-MAPK-15 (URX). Arrow, dendrite ending. (C) Schematic showing quantification of superfolderGFP-MAPK-15 enrichment at dendrite ending. Wild-type animals expressing *flp-8*pro:mCherry (URX) and *flp-8*pro:superfolderGFP-MAPK-15 (URX) were imaged. Fluorescence intensity of a 2-μm diameter circle (yellow) in a single optical plane at the dendrite ending (i, ii) or middle (iii, iv) was calculated and background fluorescence was subtracted to yield a corrected intensity (yellow numbers). Tip enrichment was calculated as the ratio of corrected intensities at the tip vs middle (i ÷ iii, ii ÷ iv). Relative enrichment of the GFP signal was calculated by first normalizing to mCherry (i ÷ ii, iii ÷ iv) to correct for local differences in cell volume, and then calculating the ratio of these normalized values ((i ÷ ii) ÷ (iii ÷ iv)). (D) Tip enrichment of superfolderGFP-MAPK-15 and mCherry. Mean ± SD: GFP, 6.3 ± 3.9, mCherry, 1.7 ± 0.7. n = 20. Colored bars, individual dendrites.

**Supplemental Figure 4.**
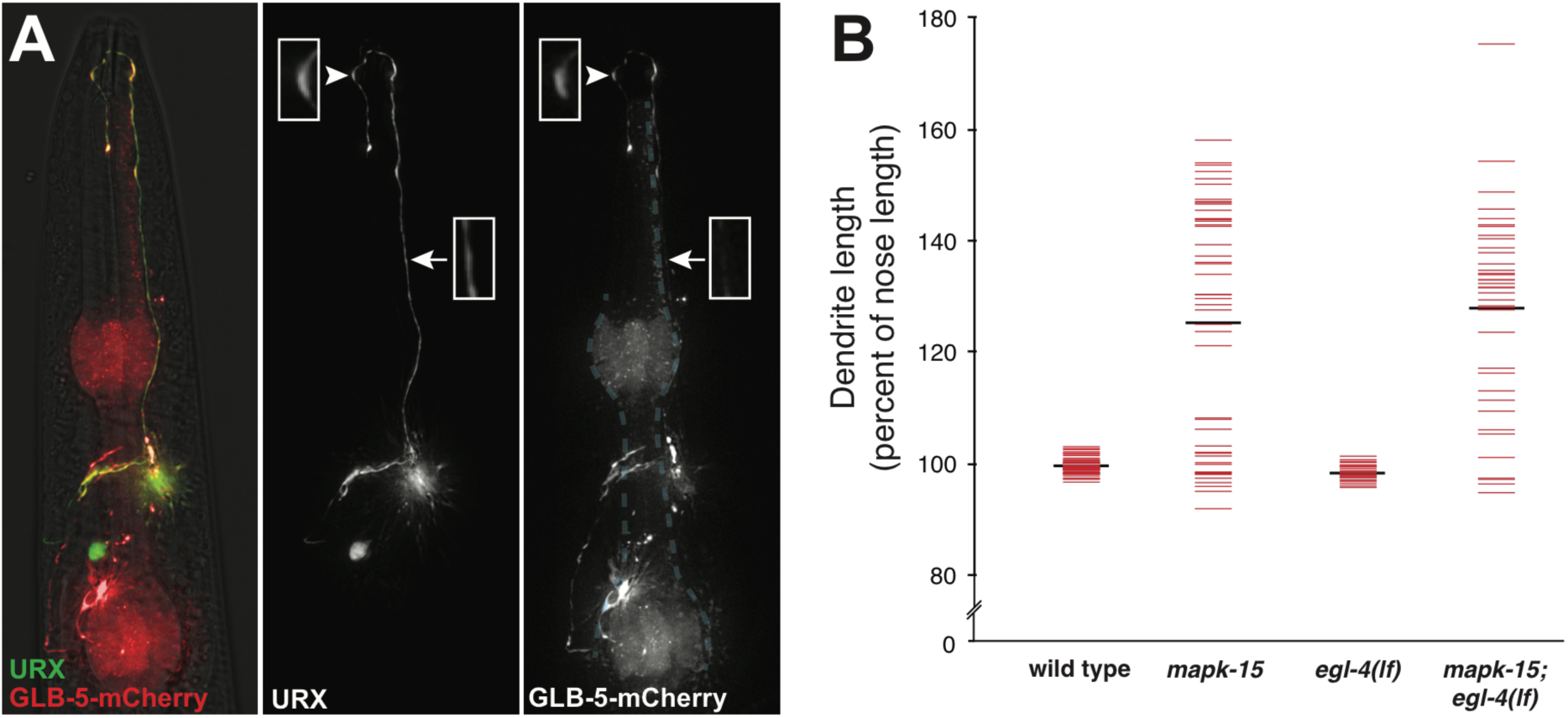
The dendritic overgrowth includes an additional sensory signaling protein and is not affected by *egl-4(lf)* (A) *mapk-15(hmn5)* animal expressing *flp-8*pro:GFP (URX) and *glb-5*pro:*glb-5*-mCherry. The *glb-5* construct is expressed in other neurons and diffusely in the pharynx (blue dashed outline). Boxes show magnifications of regions from the URX overgrowth (arrowheads) and dendrite middle (arrows). (B) Wild-type, *mapk-15*, *egl-4(lf),* and *mapk-15;egl-4(lf)* animals were synchronized as two-day adults and dendrite and nose lengths were measured. Colored bars, individual dendrites; black bars, population averages. n>33 for each genotype.

## REFERENCES

1. Berry KP, Nedivi E (2016) Experience-Dependent Structural Plasticity in the Visual System. Annu Rev Vis Sci 2: 17–35.

2. Fu AK, Ip NY (2017) Regulation of postsynaptic signaling in structural synaptic plasticity. Curr Opin Neurobiol 45: 148–155.

3. Nechipurenko IV, Doroquez DB, Sengupta P (2013) Primary cilia and dendritic spines: different but similar signaling compartments. Mol Cells 36: 288–303.

4. Mukhopadhyay S, Lu Y, Shaham S, Sengupta P (2008) Sensory signaling-dependent remodeling of olfactory cilia architecture in C. elegans. Dev Cell 14: 762–774.

5. Silverman MA, Leroux MR (2009) Intraflagellar transport and the generation of dynamic, structurally and functionally diverse cilia. Trends Cell Biol 19: 306–316.

6. Di Cristo G, Wu C, Chattopadhyaya B, Ango F, Knott G, Welker E et al. (2004) Subcellular domain-restricted GABAergic innervation in primary visual cortex in the absence of sensory and thalamic inputs. Nat Neurosci 7: 1184–1186.

7. Megías M, Emri Z, Freund TF, Gulyás AI (2001) Total number and distribution of inhibitory and excitatory synapses on hippocampal CA1 pyramidal cells. Neuroscience 102: 527–540.

8. Branco T, Häusser M (2010) The single dendritic branch as a fundamental functional unit in the nervous system. Curr Opin Neurobiol 20: 494–502.

9. White JG, Southgate E, Thomson JN, Brenner S (1986) The structure of the nervous system of the nematode Caenorhabditis elegans. Philos Trans R Soc Lond B Biol Sci 314: 1–340.

10. Ward S, Thomson N, White JG, Brenner S (1975) Electron microscopical reconstruction of the anterior sensory anatomy of the nematode Caenorhabditis elegans.?2UU. J Comp Neurol 160: 313–337.

11. Chisholm AD, Hutter H, Jin Y, Wadsworth WG (2016) The Genetics of Axon Guidance and Axon Regeneration in Caenorhabditis elegans. Genetics 204: 849–882.

12. Dong X, Liu OW, Howell AS, Shen K (2013) An extracellular adhesion molecule complex patterns dendritic branching and morphogenesis. Cell 155: 296–307.

13. Díaz-Balzac CA, Rahman M, Lázaro-Peña MI, Martin Hernandez LA, Salzberg Y, Aguirre-Chen C et al. (2016) Muscle- and Skin-Derived Cues Jointly Orchestrate Patterning of Somatosensory Dendrites. Curr Biol 26: 2379–2387.

14. Liang X, Dong X, Moerman DG, Shen K, Wang X (2015) Sarcomeres Pattern Proprioceptive Sensory Dendritic Endings through UNC-52/Perlecan in C. elegans. Dev Cell 33: 388–400.

15. Liu OW, Shen K (2011) The transmembrane LRR protein DMA-1 promotes dendrite branching and growth in C. elegans. Nat Neurosci 15: 57–63.

16. Liu X, Wang X, Shen K (2016) Receptor tyrosine phosphatase CLR-1 acts in skin cells to promote sensory dendrite outgrowth. Dev Biol 413: 60–69.

17. Oren-Suissa M, Hall DH, Treinin M, Shemer G, Podbilewicz B (2010) The fusogen EFF-1 controls sculpting of mechanosensory dendrites. Science 328: 1285–1288.

18. Salzberg Y, Díaz-Balzac CA, Ramirez-Suarez NJ, Attreed M, Tecle E, Desbois M et al. (2013) Skin-derived cues control arborization of sensory dendrites in Caenorhabditis elegans. Cell 155: 308–320.

19. Salzberg Y, Ramirez-Suarez NJ, Bülow HE (2014) The proprotein convertase KPC-1/furin controls branching and self-avoidance of sensory dendrites in Caenorhabditis elegans. PLoS Genet 10: e1004657.

20. Smith CJ, Watson JD, Spencer WC, O’Brien T, Cha B, Albeg A et al. (2010) Time-lapse imaging and cell-specific expression profiling reveal dynamic branching and molecular determinants of a multi-dendritic nociceptor in C. elegans. Dev Biol 345: 18–33.

21. Taylor CA, Yan J, Howell AS, Dong X, Shen K (2015) RAB-10 Regulates Dendritic Branching by Balancing Dendritic Transport. PLoS Genet 11: e1005695.

22. Wei X, Howell AS, Dong X, Taylor CA, Cooper RC, Zhang J et al. (2015) The unfolded protein response is required for dendrite morphogenesis. Elife 4: e06963.

23. Zou W, Shen A, Dong X, Tugizova M, Xiang YK, Shen K (2016) A multi-protein receptor-ligand complex underlies combinatorial dendrite guidance choices in C. elegans. Elife 5:

24. Smith CJ, Watson JD, VanHoven MK, Colón-Ramos DA, Miller DM (2012) Netrin (UNC-6) mediates dendritic self-avoidance. Nat Neurosci 15: 731–737.

25. Yip ZC, Heiman MG (2016) Duplication of a Single Neuron in C. elegans Reveals a Pathway for Dendrite Tiling by Mutual Repulsion. Cell Rep 15: 2109–2117.

26. Hilliard MA, Bargmann CI (2006) Wnt signals and frizzled activity orient anterior-posterior axon outgrowth in C. elegans. Dev Cell 10: 379–390.

27. Maniar TA, Kaplan M, Wang GJ, Shen K, Wei L, Shaw JE et al. (2011) UNC-33 (CRMP) and ankyrin organize microtubules and localize kinesin to polarize axon-dendrite sorting. Nat Neurosci 15: 48–56.

28. Doroquez DB, Berciu C, Anderson JR, Sengupta P, Nicastro D (2014) A high-resolution morphological and ultrastructural map of anterior sensory cilia and glia in Caenorhabditis elegans. Elife 3: e01948.

29. Perkins LA, Hedgecock EM, Thomson JN, Culotti JG (1986) Mutant sensory cilia in the nematode Caenorhabditis elegans. Dev Biol 117: 456–487.

30. Heiman MG, Shaham S (2009) DEX-1 and DYF-7 establish sensory dendrite length by anchoring dendritic tips during cell migration. Cell 137: 344–355.

31. Cheung BH, Arellano-Carbajal F, Rybicki I, de Bono M (2004) Soluble guanylate cyclases act in neurons exposed to the body fluid to promote C. elegans aggregation behavior. Curr Biol 14: 1105–1111.

32. Gray JM, Karow DS, Lu H, Chang AJ, Chang JS, Ellis RE et al. (2004) Oxygen sensation and social feeding mediated by a C. elegans guanylate cyclase homologue. Nature 430: 317–322.

33. Gross E, Soltesz Z, Oda S, Zelmanovich V, Abergel Z, de Bono M (2014) GLOBIN-5-dependent O2 responses are regulated by PDL-1/PrBP that targets prenylated soluble guanylate cyclases to dendritic endings. J Neurosci 34: 16726–16738.

34. Laurent P, Soltesz Z, Nelson GM, Chen C, Arellano-Carbajal F, Levy E et al. (2015) Decoding a neural circuit controlling global animal state in C. elegans. Elife 4:

35. Zimmer M, Gray JM, Pokala N, Chang AJ, Karow DS, Marletta MA et al. (2009) Neurons detect increases and decreases in oxygen levels using distinct guanylate cyclases. Neuron 61: 865–879.

36. Kim K, Li C (2004) Expression and regulation of an FMRFamide-related neuropeptide gene family in Caenorhabditis elegans. J Comp Neurol 475: 540–550.

37. Seidman CE, Seidman JG (1998) Molecular genetic studies of familial hypertrophic cardiomyopathy. Basic Res Cardiol 93 Suppl 3: 13–16.

38. Takashima Y, Kitaoka S, Bando T, Kagawa H (2012) Expression profiles and unc-27 mutation rescue of the striated muscle type troponin I isoform-3 in Caenorhabditis elegans. Genes Genet Syst 87: 243–251.

39. Bermingham DP, Hardaway JA, Refai O, Marks CR, Snider SL, Sturgeon SM et al. (2017) The Atypical MAP Kinase SWIP-13/ERK8 Regulates Dopamine Transporters through a Rho-Dependent Mechanism. J Neurosci 37: 9288–9304.

40. Kazatskaya A, Kuhns S, Lambacher NJ, Kennedy JE, Brear AG, McManus GJ et al. (2017) Primary Cilium Formation and Ciliary Protein Trafficking Is Regulated by the Atypical MAP Kinase MAPK15 in Caenorhabditis elegans and Human Cells. Genetics

41. Piasecki BP, Sasani TA, Lessenger AT, Huth N, Farrell S (2017) MAPK-15 is a ciliary protein required for PKD-2 localization and male mating behavior in Caenorhabditis elegans. Cytoskeleton (Hoboken) 74: 390–402.

42. Brenner S (1974) The genetics of Caenorhabditis elegans. Genetics 77: 71–94.

43. McKeown C, Praitis V, Austin J (1998) sma-1 encodes a betaH-spectrin homolog required for Caenorhabditis elegans morphogenesis. Development 125: 2087–2098.

44. Praitis V, Ciccone E, Austin J (2005) SMA-1 spectrin has essential roles in epithelial cell sheet morphogenesis in C. elegans. Dev Biol 283: 157–170.

45. Shaham S (2010) Chemosensory organs as models of neuronal synapses. Nat Rev Neurosci 11: 212–217.

46. Abe MK, Saelzler MP, Espinosa R, Kahle KT, Hershenson MB, Le Beau MM et al. (2002) ERK8, a new member of the mitogen-activated protein kinase family. J Biol Chem 277: 16733–16743.

47. Klevernic IV, Stafford MJ, Morrice N, Peggie M, Morton S, Cohen P (2006) Characterization of the reversible phosphorylation and activation of ERK8. Biochem J 394: 365–373.

48. Colecchia D, Strambi A, Sanzone S, Iavarone C, Rossi M, Dall’Armi C et al. (2012) MAPK15/ERK8 stimulates autophagy by interacting with LC3 and GABARAP proteins. Autophagy 8: 1724–1740.

49. Colecchia D, Rossi M, Sasdelli F, Sanzone S, Strambi A, Chiariello M (2015) MAPK15 mediates BCR-ABL1-induced autophagy and regulates oncogene-dependent cell proliferation and tumor formation. Autophagy 11: 1790–1802.

50. Johnson JL, Leroux MR (2010) cAMP and cGMP signaling: sensory systems with prokaryotic roots adopted by eukaryotic cilia. Trends Cell Biol 20: 435–444.

51. Raizen DM, Cullison KM, Pack AI, Sundaram MV (2006) A novel gain-of-function mutant of the cyclic GMP-dependent protein kinase egl-4 affects multiple physiological processes in Caenorhabditis elegans. Genetics 173: 177–187.

52. Busch KE, Laurent P, Soltesz Z, Murphy RJ, Faivre O, Hedwig B et al. (2012) Tonic signaling from O_2_ sensors sets neural circuit activity and behavioral state. Nat Neurosci 15: 581–591.

53. Cheung BH, Cohen M, Rogers C, Albayram O, de Bono M (2005) Experience-dependent modulation of C. elegans behavior by ambient oxygen. Curr Biol 15: 905–917.

54. McGrath PT, Rockman MV, Zimmer M, Jang H, Macosko EZ, Kruglyak L et al. (2009) Quantitative mapping of a digenic behavioral trait implicates globin variation in C. elegans sensory behaviors. Neuron 61: 692–699.

55. Weber KP, De S, Kozarewa I, Turner DJ, Babu MM, de Bono M (2010) Whole genome sequencing highlights genetic changes associated with laboratory domestication of C. elegans. PLoS One 5: e13922.

56. Marshall WF (2015) How Cells Measure Length on Subcellular Scales. Trends Cell Biol 25: 760–768.

57. Dickstein DL, Weaver CM, Luebke JI, Hof PR (2013) Dendritic spine changes associated with normal aging. Neuroscience 251: 21–32.

58. Scheibel ME, Lindsay RD, Tomiyasu U, Scheibel AB (1975) Progressive dendritic changes in aging human cortex. Exp Neurol 47: 392–403.

59. Vaughan DW (1977) Age-related deterioration of pyramidal cell basal dendrites in rat auditory cortex. J Comp Neurol 171: 501–515.

60. Buell SJ, Coleman PD (1979) Dendritic growth in the aged human brain and failure of growth in senile dementia. Science 206: 854–856.

61. Connor JR, Diamond MC, Connor JA, Johnson RE (1981) A Golgi study of dendritic morphology in the occipital cortex of socially reared aged rats. Exp Neurol 73: 525–533.

62. Lee WC, Huang H, Feng G, Sanes JR, Brown EN, So PT et al. (2006) Dynamic remodeling of dendritic arbors in GABAergic interneurons of adult visual cortex. PLoS Biol 4: e29.

63. Schroeder NE, Androwski RJ, Rashid A, Lee H, Lee J, Barr MM (2013) Dauer-specific dendrite arborization in C. elegans is regulated by KPC-1/Furin. Curr Biol 23: 1527–1535.

64. Kravtsov V, Oren-Suissa M, Podbilewicz B (2017) The fusogen AFF-1 can rejuvenate the regenerative potential of adult dendritic trees by self-fusion. Development 144: 2364–2374.

65. Yu S, Avery L, Baude E, Garbers DL (1997) Guanylyl cyclase expression in specific sensory neurons: a new family of chemosensory receptors. Proc Natl Acad Sci U S A 94: 3384–3387.

66. Pédelacq JD, Cabantous S, Tran T, Terwilliger TC, Waldo GS (2006) Engineering and characterization of a superfolder green fluorescent protein. Nat Biotechnol 24: 79–88.

67. Kodama-Namba E, Fenk LA, Bretscher AJ, Gross E, Busch KE, de Bono M (2013) Cross-modulation of homeostatic responses to temperature, oxygen and carbon dioxide in C. elegans. PLoS Genet 9: e1004011.

68. Doitsidou M, Poole RJ, Sarin S, Bigelow H, Hobert O (2010) C. elegans mutant identification with a one-step whole-genome-sequencing and SNP mapping strategy. PLoS One 5: e15435.

69. Minevich G, Park DS, Blankenberg D, Poole RJ, Hobert O (2012) CloudMap: a cloud-based pipeline for analysis of mutant genome sequences. Genetics 192: 1249–1269.

